# Pathway-Level Integration of Proteogenomic Data in Breast Cancer Using Independent Component Analysis

**DOI:** 10.1101/175687

**Authors:** Wenke Liu, Sisi Ma, David Fenyö

## Abstract

Recent advances in the multi-omics characterization necessitate pathway-level abstraction and knowledge integration across different data types. In this study, we apply independent component analysis (ICA) to human breast cancer proteogenomics data to retrieve mechanistic information. We show that as an unsupervised feature extraction method, ICA was able to construct signatures with known biological relevance on both transcriptome and proteome levels. Moreover, proteome and transcriptome signatures can be associated by their respective correlation with patient clinical features, providing an integrated description of phenotype-related biological processes. Our results demonstrate that the application of ICA to proteogenomics data could lead to pathway-level knowledge discovery. Potential extension of this approach to other data and cancer types may contribute to pan-cancer integration of multi-omics information.

## INTRODUCTION

Breast cancer is the most common cancer among women, and while targeted therapies have helped to significantly reduced breast cancer mortality rate in the past decade, further improvement will require a comprehensive understanding of the molecular mechanisms of the disease^1,2^. Recently, deep mass spectrometry based proteomic characterization of genomically annotated breast cancer samples by the Clinical Proteomic Tumor Analysis Consortium (CPTAC) has marked the initial step of an proteogenomic integrative approach, in which recurrent mutations and copy number variations on the genomic level, expression profiles on the transcriptomic level and protein abundance and functional manifestations on proteomic level were measured for the same group of patient samples and examined in the same framework^3–5^. The collection of high quality multi-omics data immediately led to the discovery of concordant gene amplification and protein phosphorylation in key pathways^3^. At the same time, there is increasing demand for integrative analysis that could incorporate all data types and extract pathway level. Since in all human patients ‘-omics’ data sets the number of features far exceeds the number of samples, analysis of any single data type is already susceptible to ‘the curse of dimensionality’, and integration by simple concatenation of multi-omics data would be an even less desirable option. Our previous work has benchmarked the predictive power of multi-omics datasets for classifying breast cancer patients into different survival groups and showed that combined multi-omics datasets produced with data-driven fusion techniques were not able to outperform proteomic data alone^6^. This result highlighted the possible redundancy among information contained in different biological levels, and motivates us to explore other data fusion techniques that extract both concordant and complementary features from high-dimensional multi-omics data.

In the current study, we applied independent component analysis to proteomic and transcriptomic data of 77 breast cancer samples to extract pathway-level molecular signatures. Independent component analysis (ICA) is an unsupervised learning method widely used in signal processing and has been applied to cancer genomics with notable success^7–9^. This approach decomposes the molecular profiles into linear combinations of non-Gaussian independent sources or components, each of which is comprised of weighted contributions from individual genes. Therefore, ICA reduces the dimensionality of original data by representing the molecular profile of each sample as weighted sum of several ‘meta-genes’ or ‘meta-proteins’, and the weight of specific meta-gene/protein (mixing scores) in one sample reflects the ‘activity’ of that component in the sample. As clinical features are also available for the CPTAC samples, molecular signatures can be constructed from clusters of meta-genes/proteins that show activity patterns correlated with these clinical features. Furthermore, taking advantage of a specific clinical feature as a ‘anchor’, this method may help extract patterns at different biological levels, which may originate from the same cellular functionality (Figure 1). The signatures extracted from different data sets were filtered based on their intrinsic stability and association with known clinical features (see Methods), and grouped into modules that showed similar correlation patterns to clinical features. Subsequent gene set enrichment analysis revealed the biological relevance of these modules to pathways such as HER2 signaling, mitosis and histone modification. Our analysis has demonstrated that ICA was able to blindly extract biologically meaningful information at pathway level. With input from clinical features or other sample sub-grouping indices, these signatures may be further integrated into multi-level models that provide insights into the molecular mechanisms of breast cancer.

**Figure 1.**
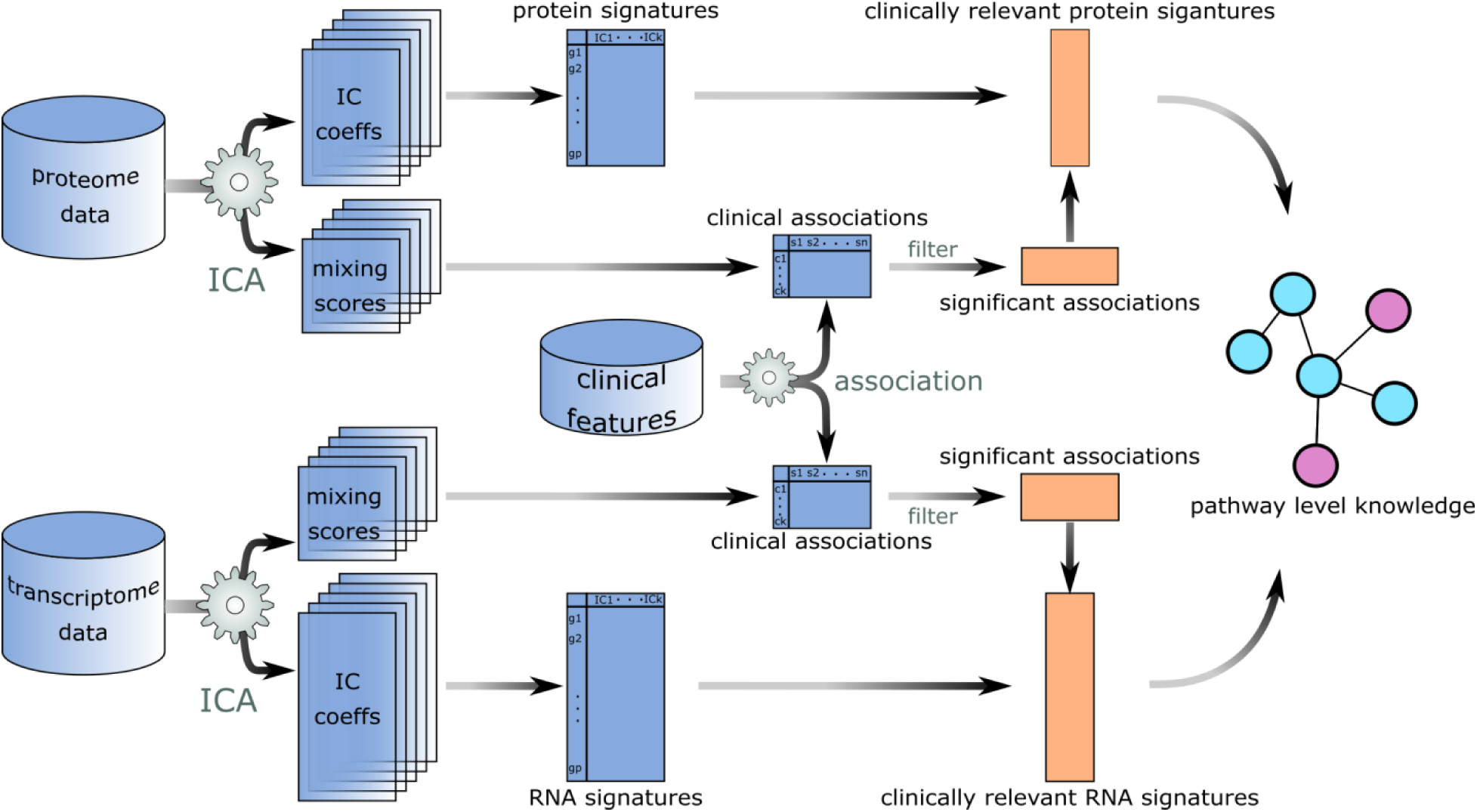
Data analysis work flow. Coefficients of independent components and their corresponding mixing scores are extracted from both proteome and transcriptome data sets for multiple randomly initiated runs. Each independent component is a pathway-level representation. Molecular signatures were identified as cluster centroids of these components, and clinical features were used to select biologically relevant signatures, which were further annotated with pathway analysis.

## RESULTS

### Stable molecular signatures extracted from proteome and transcriptome data

For both the proteome and transcriptome datasets we identified 77 clusters of meta-proteins or meta-genes from independent components obtained from 50 randomly initiated runs (Method). Centers of the stable meta-gene and meta-protein clusters, which could be found by averaging gene coefficients within each cluster, could represent a pathway-level signature. The stability of these signatures could be inspected by visualizing all meta-genes and meta-proteins with t-Distributed Stochastic Neighbor Embedding (t-SNE), a widely used dimensionality reduction technique^10^ (Figure 2a, c). A large portion of the meta-gene and meta-protein clusters are compact, while the rest formed a visually indistinguishable mixture, consistent with the observation that the average silhouette width of all clusters followed a bimodal distribution (Figure 2b, d). Another metric that could help evaluate the stability of extracted signatures is the number of different runs that the members of each cluster originated from. It has been proposed that reliable independent components should be highly recurrent, therefore, a cluster the members of which were extracted from different runs would be considered as more stable. Indeed, this metric largely agrees with silhouette width, such that a lot of compact clusters were also comprised of meta-proteins or meta-genes extracted from all 50 runs.

**Figure 2.**
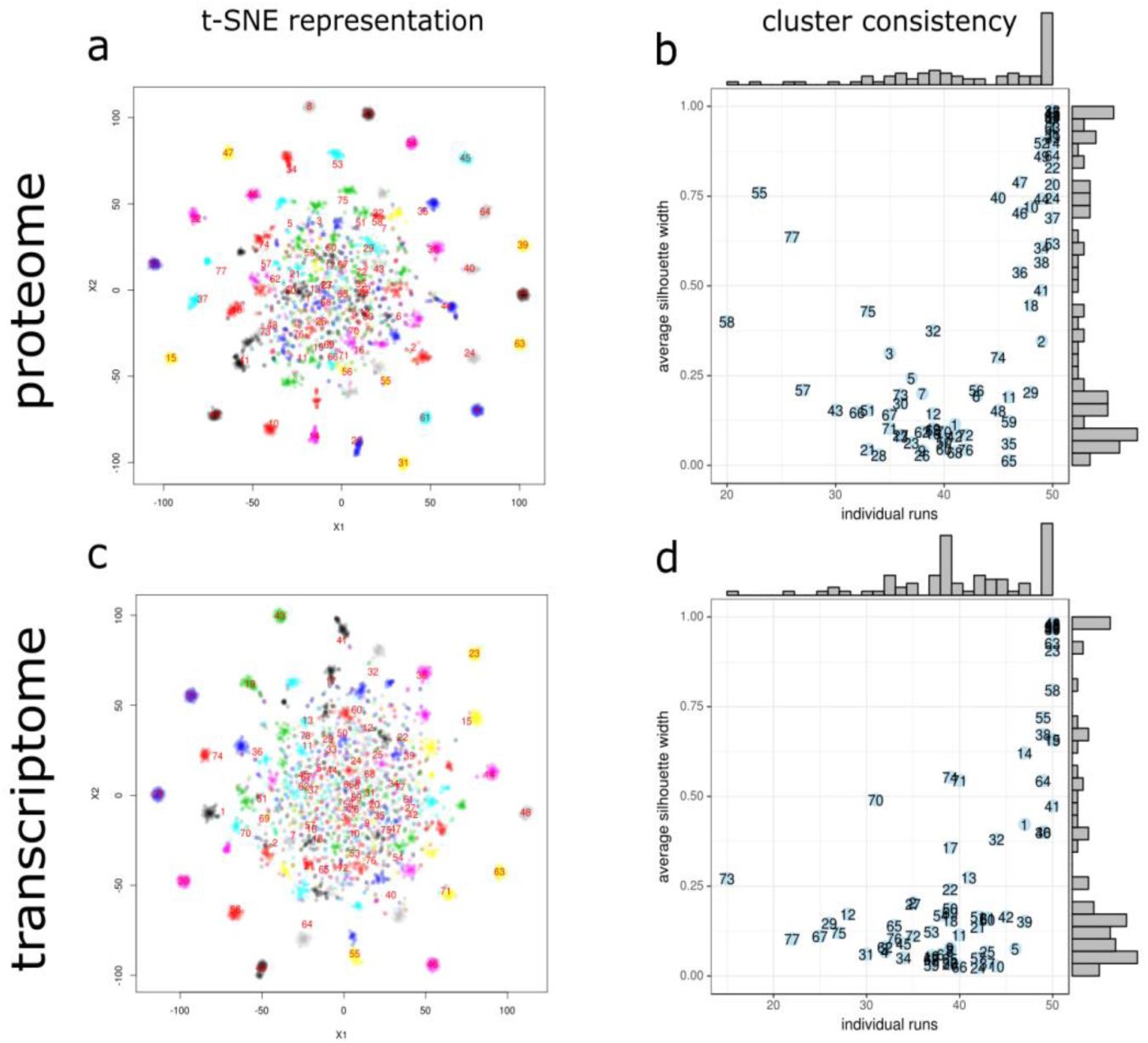
Evaluation of component consistency. **a**, **c**, t-SNE representation of components obtained from 50 randomly initiated runs. Colors represent cluster assignments. **b**, **d**, average silhouette widths for the 77 clusters were plotted against the number of different individual runs from which the members of each cluster were extracted. Marginal histograms revealed the distribution of the metrics. Large number indicates that the cluster is recurrent and more likely to capture true structures in the data.

### Mixing scores reveal clinical relevance of identified components

In addition to their numerical stability, the clusters of components representing groups of meta-proteins or meta-genes identified from different ICA runs can also be evaluated with their relevance to known characteristics of breast cancer. For each particular ICA decomposition, the rows of the mixing score matrices represented the ‘activity level’ of the corresponding independent components in all of the samples. Therefore, it is possible to establish associations between the activity patterns of meta-genes and meta-proteins and clinical features available for TCGA samples, which would in turn reveal the functional relevance of the signatures. We recoded 22 clinical features into ordinal factors and use linear regression to assess their correlation with activity scores of meta-genes or meta-proteins in each signature cluster. A stringent threshold of nominal *P*-value (10^-5^) was used to correct for multiple comparisons and select the most significant associations. Within each cluster, the number of meta-proteins or meta-genes with activity levels that showed significant linear relation with the 22 clinical features was documented. Large number of significant association between a signature cluster and a clinical feature indicates that the signature may contain pathway-level information about molecular mechanisms underlying the clinical feature. 33 out of the 77 proteome IC clusters showed significant association with 10 clinical features, while 60 transcriptome clusters were correlated with 12 clinical features (Figure 3).

**Figure 3.**
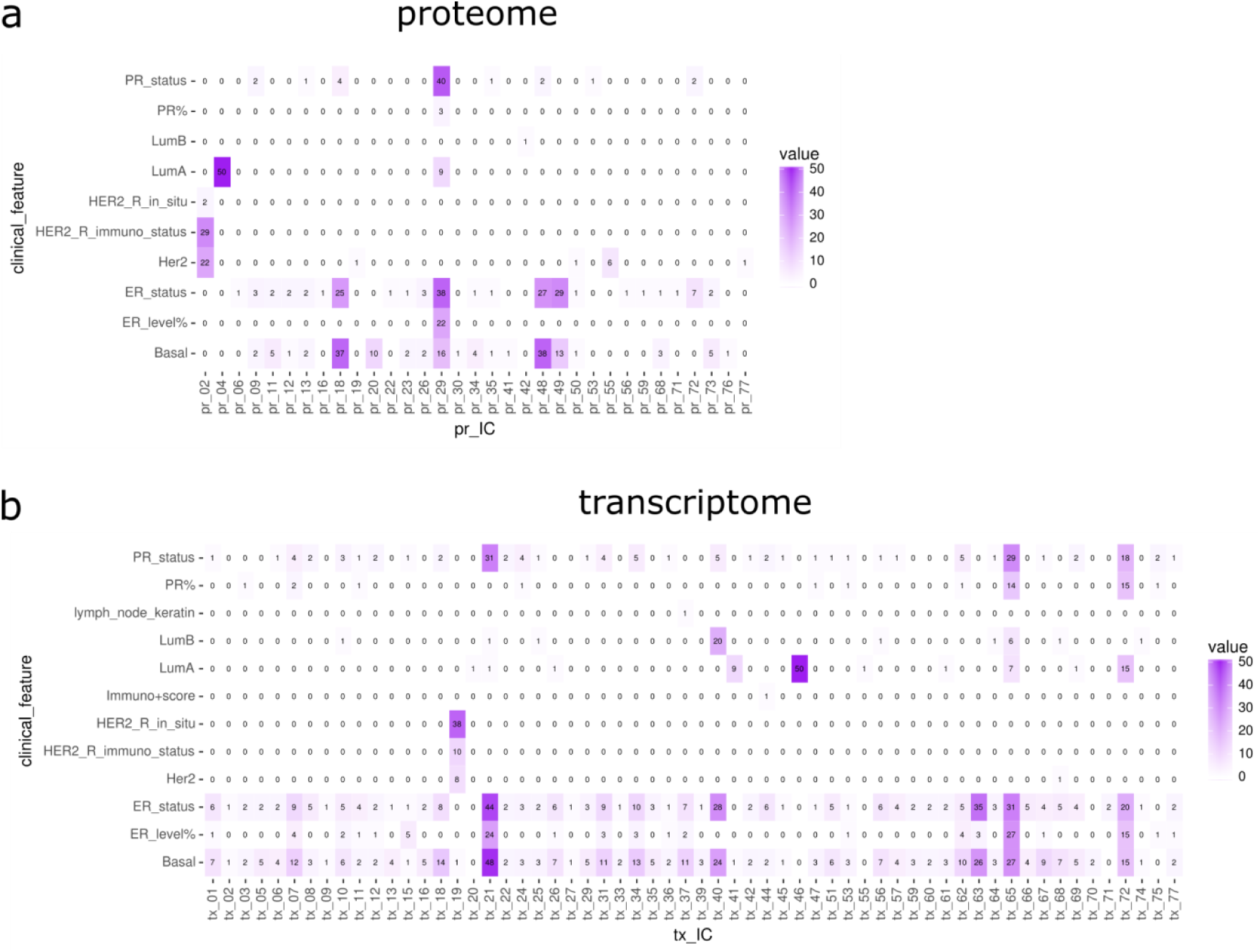
Clinical associations of IC clusters. Number of significant associations found between corresponding mixing scores and clinical features for ICs in each cluster. Only non-zero columns and rows were shown.

The biological relevance of the identified clusters could be further validated by inspecting the average gene coefficients (the centroids of the clusters). For example, the proteome signature cluster that contains 29 members that were significantly correlated with the HER2 receptor immuno status and 22 members associated with the Her2 subtype index (pr_02), was heavily weighted on ERBB4 and ERBB2, two tyrosine receptor kinases that mediate the Her2 signaling, and the two proteins were assigned the two largest coefficients (11.13 and 10.68, Table 1). Another signature (pr_29) correlated with estrogen receptor (ER) and progesterone receptor (PR) status also exhibited high average scores for PGR (7.73) and ESR1 (4.42) proteins. Interestingly, the signature pr_04, which displayed strong correlation with the LumA subtype, is enriched with a group of mitosis-related proteins, indicating that cell division checkpoint may be differentially regulated in LumA subtype.

**Table 1:**
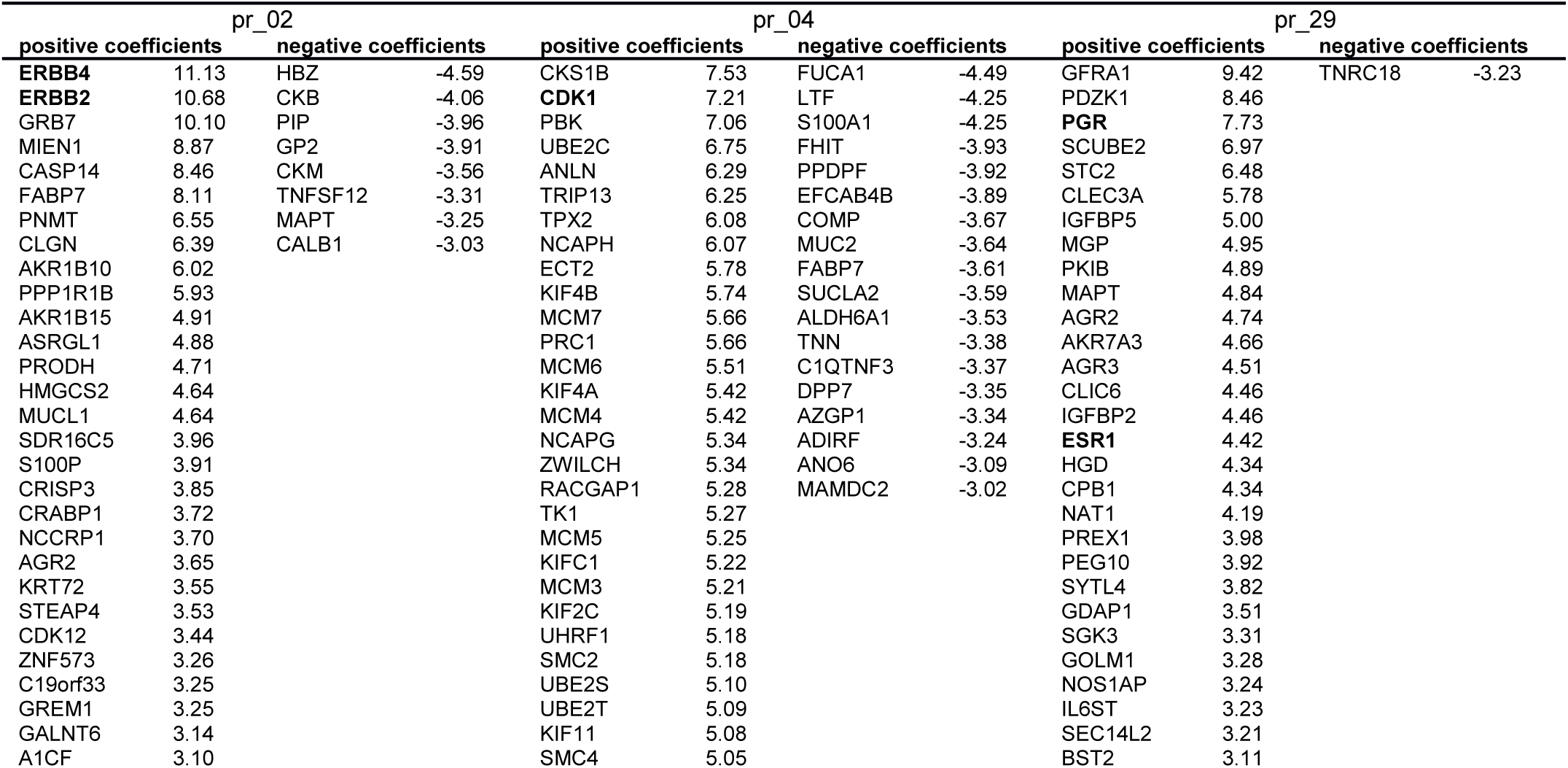
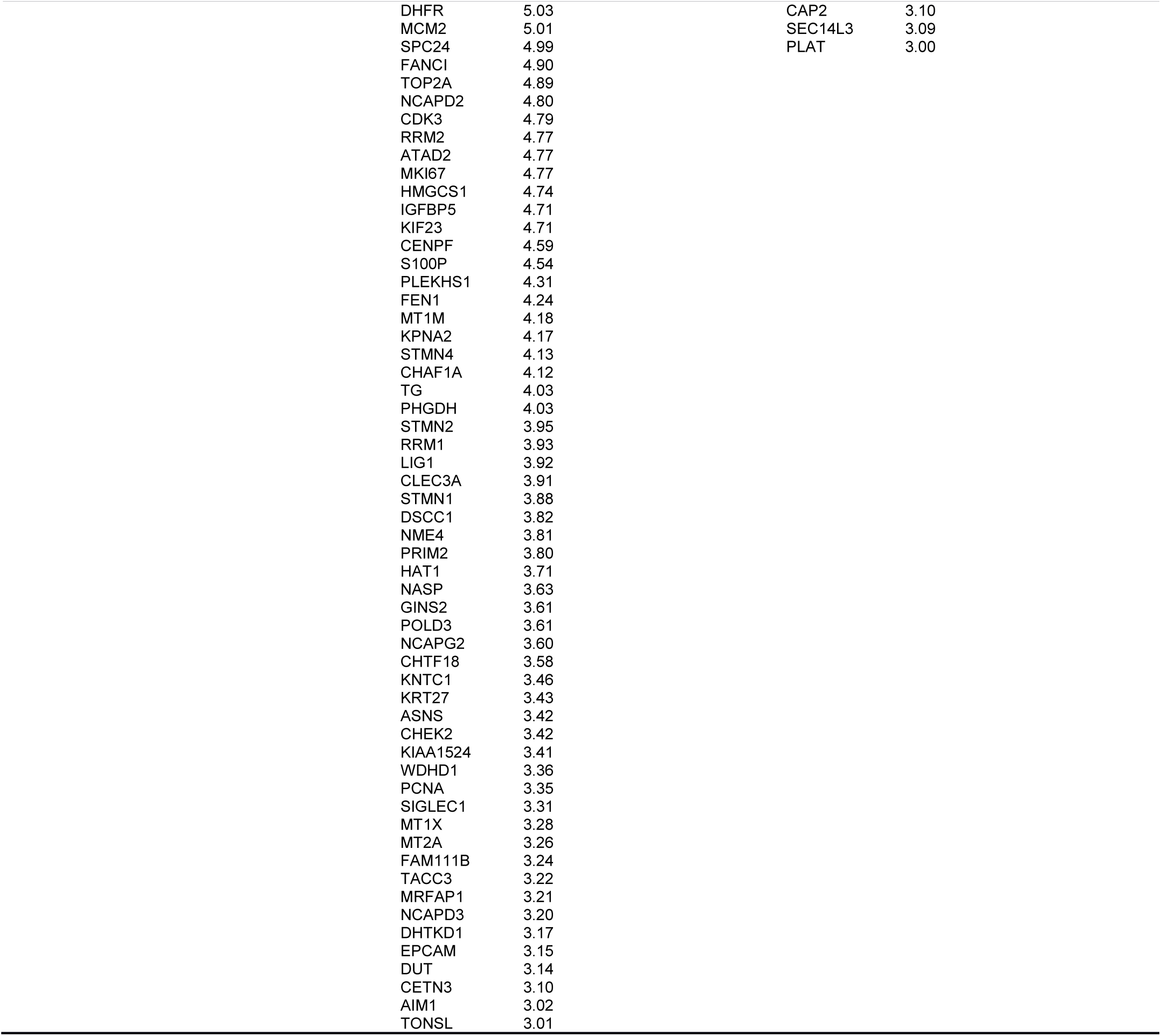
Average coefficients of selected proteome signatures.

Large value coefficients of known marker genes revealed the biological relevance of a subset of the molecular signatures, but more biological information could be exploited on the pathway level. The coefficient vector for each signature is a pre-ranked gene list that can be subjected to Gene Set Enrichment Analysis (GSEA) on curated gene sets and GO terms^11^, and the collective effect of genes with coefficients of smaller absolute values could contribute to the enrichment metric. GSEA results showed that the clinical-related proteome IC clusters retrieved a lot of the breast cancer related gene sets determined by experimental manipulations, as well as gene sets that characterized basic biological processes (Table 2). For example, the proteome signatures pr_02 and pr_29, which were strongly associated with Her2 and ER status, also exhibited enrichment of the corresponding gene sets (Table 2). The transcriptome signature tx_07, which were correlated with Basal subtype, ER and PR status, showed negative enrichment of targets of LSD1, a histone demethylase of H3K4 sites, which had known links to be poor breast cancer prognostics and ER negative status^12,13^. Interestingly, the same signature cluster also exhibited positive enrichment of genes with H3K27Me3 sites, indicating that epigenetic regulation is highly orchestrated in breast cancer.

**Table 2:**
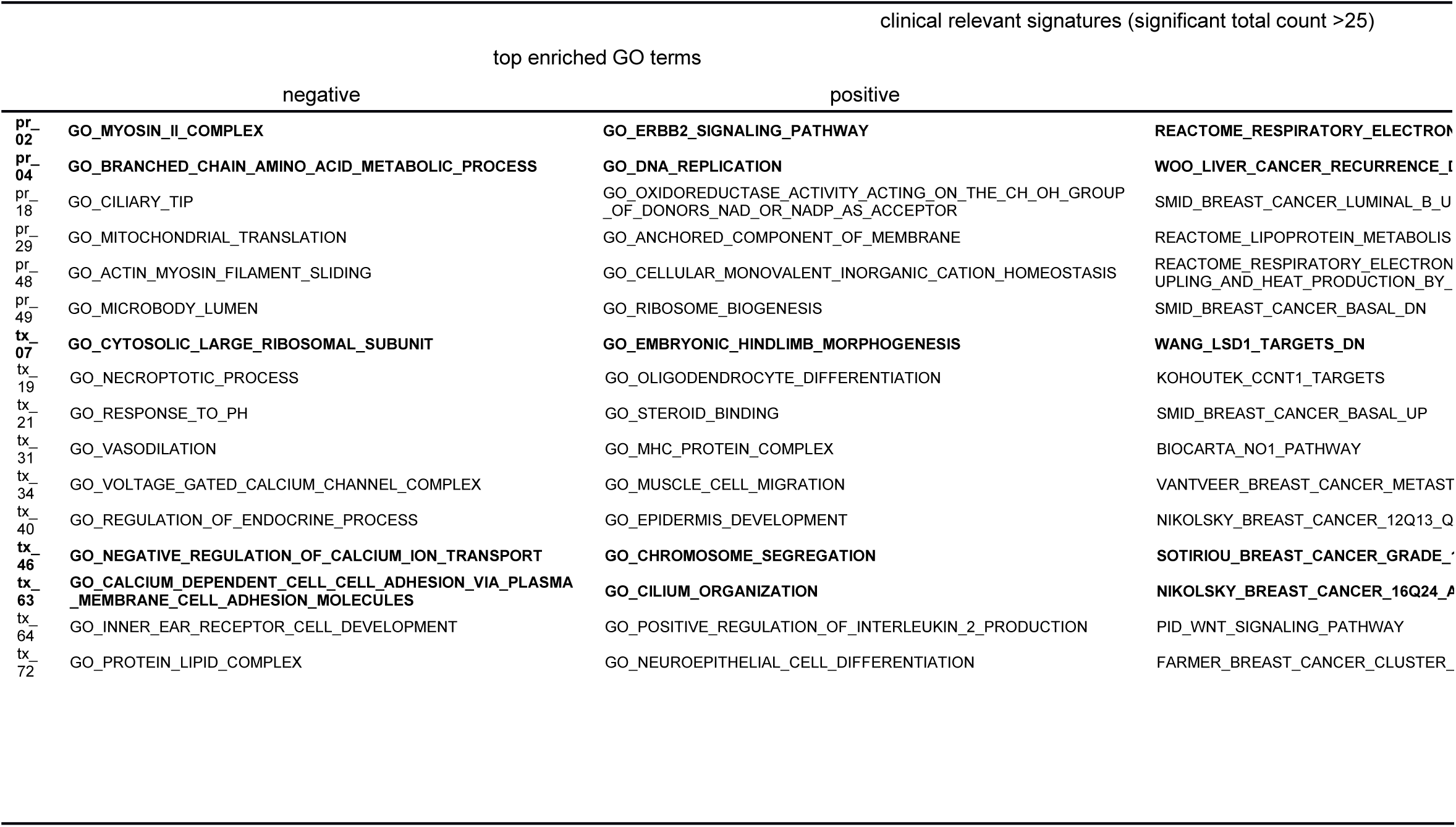
GO terms and curated gene sets enrichment of selected signatures.

Although the most clinically-relevant clusters (total count of significant association >25) contained proteome and transcriptome signatures that are highly recurrent, these clusters were not very homogeneous (Supplementary table 1). On the other hand, out of the 20 most consistent clusters with average silhouette widths larger than 0.9 and are recurrent in every run for both proteome and transcriptome, only 3 signatures (pr_04, tx_46, tx_63) showed direct clinical relevance (Supplementary table 1). The meta-proteins and meta-genes in the clusters of pr_04 and tx_46 were both significantly associated with Luminal A subtype index, and showed enrichment of genes involved in estrogen response (table 2). The signature tx_63 was associated with the Basal subtype and ER status, and its coefficients exhibited significant negative enrichment of genes in 16q24 loci (*P*<0.001, FDR<0.001), a breast cancer risk factor, as well as positive enrichment of genes involved in endocrine therapy response (*P*<0.001, FDR<0.001). The large extent of disagreement between clinical relevance and cluster consistency suggest that ICA may serve as a promising approach to knowledge discovery, as stable clusters that did not show clinical relevance may provide new insights into the molecular mechanisms of breast cancer.

### Integrative pathway analysis of proteomic and transcriptomic components guided by clinical features

The correlation between mixing scores and clinical features provided a valuable opportunity to find the link between independently extracted components from different data types. For example, in both proteome and transcriptome analyses, there was a highly consistent signature (pr_04 and tx_46), such that meta-proteins and meta-genes in the two clusters were recurrent in all 50 randomly initiated runs and the corresponding mixing scores all showed significant correlation with index of Luminal A subtype (Figures 2 and 3). To integrate the proteomic and transcriptomic information with guidance from clinical relevance, we applied the hierarchical clustering algorithm to the vectors of clinical association counts for the most clinically relevant meta-protein and meta-gene clusters. Proteome and transcriptome signatures could therefore be grouped based on their similarity in functional indications, as the metric of direct correlation between gene coefficients of different signatures was limited by noise introduced by high dimensionality (Supplementary Figure 1). Combined network visualization of GO terms enriched in proteome and transcriptome signatures allowed us to examine complementary functional modules on different biological levels. For example, the pr_04 and tx_46 clusters may characterize Luminal A subtype specific processes from protein and RNA perspectives (Figure 4b). While majority of the gene set networks consisted of nodes and edges between proteome and transcriptome, specific networks could be identified for both data modes, with tx_46 showed specific enrichment in a DNA replication-related network, and pr_04 showed specific enrichment in polysaccharide metabolism (Figure 4c).

**Figure 4.**
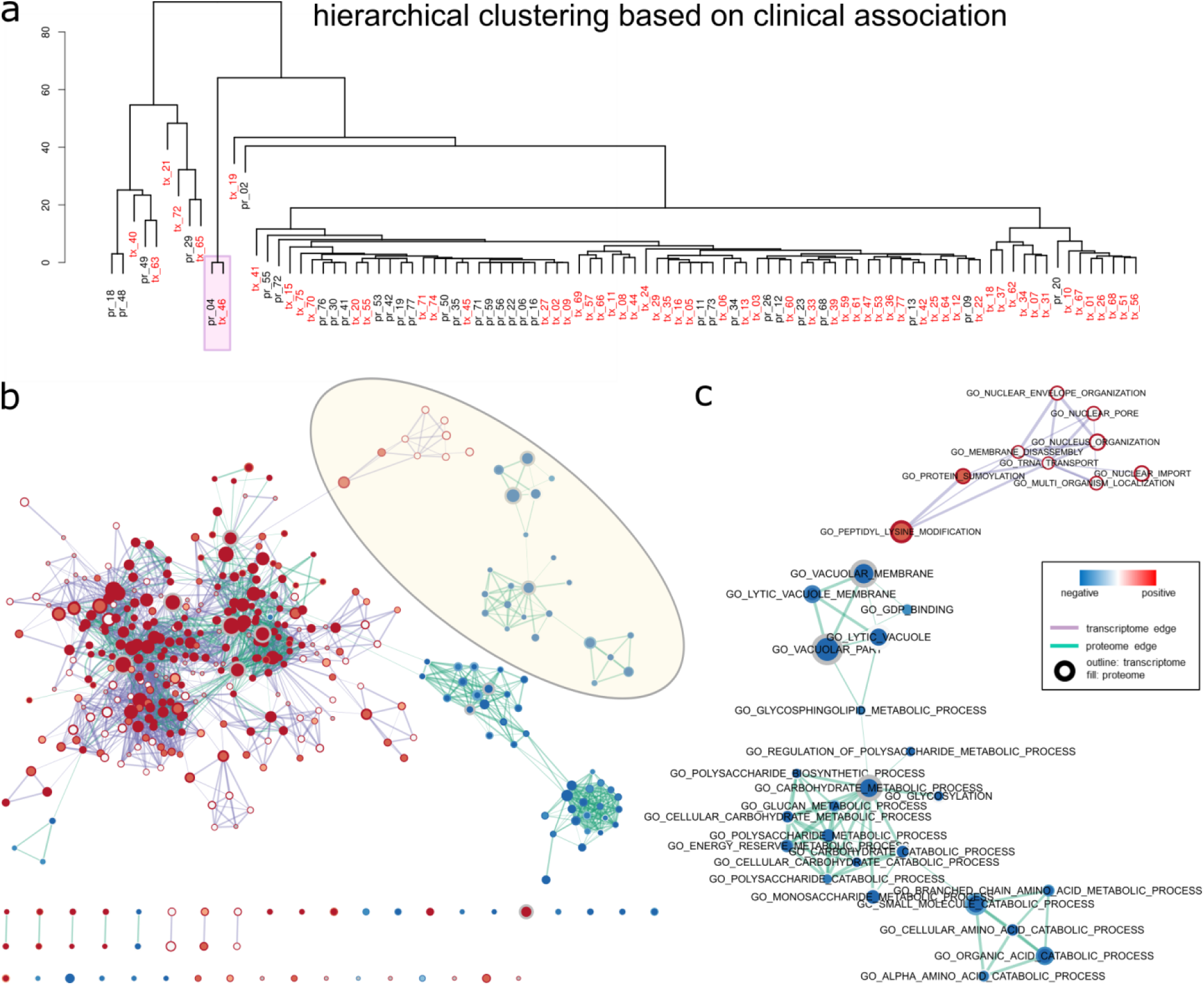
Integrative analysis of proteome and transcriptome signatures. Only clinically relevant IC clusters are shown (total count of significant associations >0). **a**, hierarchical clustering of the clinical association of all signatures. **b**, GSEA enrichment map of the combined network of cluster centers of pr_04 and tx_46 indicated in **a**. **c**, zoomed-in view of a transcriptome-specific (top) and a proteome specific subnetwork highlighted in **b**.

## DISCUSSION

We have utilized independent component analysis to gain pathway-level insights into the mechanisms of breast cancer and develop protein/gene modules as clinical signatures. Meta-proteins and meta-genes are extracted from the data in an unsupervised manner, and signatures are further selected based on consistency of meta-gene/protein clusters or the association between their activity scores and known clinical features. Gene set annotation revealed that several selected signatures contained biologically relevant information. A proteome signature (pr_02) characterizing strong activation of the Her2 pathway was recovered as a Her2-related meta-protein cluster. On the transcriptome level, meta-genes enriched for genes in the 16q24 risk loci formed a stable signature that showed correlation with both ER status and Basal subtype. Stable signatures on both proteome and transcriptome levels (pr_04 and tx_46) were found to be heavily associated with the LumA index, suggesting that cell division and growth is specifically regulated in this subtype.

As an unsupervised blind source separation method, independent component analysis has been applied to multiple biological data types. Consistent with previous reports, our results demonstrated that ICA is able to extract biological meaningful information solely based on the intrinsic structures of the transcriptome and proteome data. The success of the method suggested that ICA have captured some aspects of the processes underlying proteome and transcriptome profiles. It is reasonable to assume that the observed RNA and protein levels are the sum of several up-regulation and down-regulation modules, in which the distribution of individual gene levels deviate radically from the normal distribution that characterizes a noisy background.

Using clinical association or signature stability as selection criterion gave rise to two lists of potential signatures with only a few overlapping members (pr_04, tx_46, tx_63), suggesting that the most clinically relevant signatures are not very stable under the current method. Other clustering method such as density based clustering may be used to improve the estimation of stable signatures^14,15^. On the other hand, it is possible that the most stable signatures described the housekeeping processes common in all samples, but they may also help reveal novel molecular mechanisms of breast cancer that are not previously linked to any phenotype. As a feature construction procedure, ICA could also facilitate knowledge integration from multiple data types. In addition to the integration of proteome and transcriptome signatures as demonstrated in this work, future studies could further apply ICA on omics data sets of multiple cancer types. Extracted molecular signatures may be grouped in another round of clustering procedure to reveal pan-cancer and cancer type specific mechanisms.

## METHODS

### DATA SETS

Transcriptome and proteome data of 77 breast cancer patients were collected as part of the TCGA and CPTAC projects^3,16^. Transcriptome was characterized with the Agilent mRNA expression array platform and global protein abundance data were obtained with iTRAQ 4-plex LC-MS/MS technique^3,16^. For the proteome data, peptides mapped to the same gene (in total) were collapsed by taking the mean value. Expression levels of genes and abundance of proteins were included in the transcriptome and proteome matrices. Clinical and demographic features were associated with each tumor sample.

### INDEPENDENT COMPONENT ANALYSIS

Data were presented in a *p*×*n* matrix **X** with genes in rows and samples in columns. The goal of ICA is to decompose the *p*×*n* data matrix into the product of a source matrix **S** (*p*×*k*) and a mixing matrix **A** (*k*×*n*). The *i*th column of the source matrix represents coefficients of each of the *p* genes for the *i*th independent component. The coefficient vector of each component could be considered as *p* random samples that revealed probability distribution of a specific random variable. The mutual information between any pair of such variables is minimized, and the components are therefore statistically independent. Genes with coefficients of positive or negative values in the same component indicated that their levels may be enhanced or suppressed in the same biological process. The *i*th row of the mixing matrix represents contributions of the *i*th component in the source matrix to profiles of each of the *n* samples. Rows of the mixing matrix therefore record the activity of each components (or meta-genes/meta-proteins) across *n* samples.

The stochastic nature of most ICA algorithms entails that different randomly initiated runs would give rise to different results, and there is no guarantee that the true structure of the data could be correctly estimated from any single run^14,17^. To assess the statistical reliability of ICA results, table components were filtered based on an approach adapted from Engreitz et al., which cluster components obtained from different runs with K-medoids method^8^. In brief, ICA was run with *k*=*n* for 50 times to extract as much information as possible. For each individual component, the larger tail was designated as positive. All 50×*n* components were then considered as data points in *p* dimensional space and subjected to K-medoids clustering with Spearman correlation as the dissimilarity measure. For each cluster, the number of different runs that its members were extracted from was documented as a measure for cluster consistency, alongside with the average silhouette width. In general, clusters that with members appeared in more than 50% of all runs (25) were considered as likely to contain true biological signals (see component annotation section below).

All computations were carried out on the R platform. Package ‘fastICA’ which implements the iterative FastICA algorithm^18^ was used to extract non-Gaussian independent components with logcosh as contrast function. Components were subsequently assigned to 77 clusters using the ‘cluster’ package. Clusters were visualized with 2d t-SNE using the R package ‘tsne’.

### SIGNATURE ANNOTATION

Component clusters were annotated with GO terms by running Gene Set Enrichment Analysis against centroid coefficients as the pre-ranked gene lists^11,19^. Enrichment map of components were visualized with Cytoscape 3^20^. Each cluster was also associated with clinical features as following: First, 22 clinical features were recoded into ordinal variables (supplementary table). Second, ordinary linear regression models were built with corresponding mixing scores for members in a component cluster to predict each of the ordinal responses. Counts of significant associations between components and clinical features (*P*-value for the slope coefficient < 10^-5^) were tabulated. Hierarchical clustering with complete linkage was applied to the clinical associations of independent components clusters extracted from both transcriptome and proteome data.

### DATA AVAILABILITY

All data analyzed during this study are included in a published article^3^ and its supplementary information files.

## ACKNOWLEDGEMENTS

We would like to acknowledge funding by the National Cancer Institute (NCI) through CPTAC award U24 CA210972 and a contract 13XS068 from Leidos Biomedical Research, Inc., and by a grant from the Shifrin-Myers Breast Cancer Discovery Fund.

## AUTHOR CONTRIBUTIONS

W.L., S.M. and D.F. conceived the method. W.L. performed the analysis. The manuscript was co-written by W.L., S.M. and D.F.

## References

1. Narod, S. A., Iqbal, J. & Miller, A. B. Why have breast cancer mortality rates declined? J. Cancer Policy 5, 8–17 (2015).

2. Mendes, D. et al. The benefit of HER2-targeted therapies on overall survival of patients with metastatic HER2-positive breast cancer – a systematic review. Breast Cancer Res. 17, 140 (2015).

3. Mertins, P. et al. Proteogenomics connects somatic mutations to signalling in breast cancer. Nature 534, 55–62 (2016).

4. Zhang, H. et al. Integrated Proteogenomic Characterization of Human High-Grade Serous Ovarian Cancer. Cell 166, 755–65 (2016).

5. Zhang, B. et al. Proteogenomic characterization of human colon and rectal cancer. Nature 513, 382–7 (2014).

6. Ma, S., Ren, J. & Fenyö, D. Breast Cancer Prognostics Using Multi-Omics Data. AMIA Jt. Summits Transl. Sci. Proc. 2016, 52–9 (2016).

7. Frigyesi, A., Veerla, S., Lindgren, D. & Höglund, M. Independent component analysis reveals new and biologically significant structures in micro array data. BMC Bioinformatics 7, 290 (2006).

8. Engreitz, J. M., Daigle, B. J., Marshall, J. J. & Altman, R. B. Independent component analysis: mining microarray data for fundamental human gene expression modules. J. Biomed. Inform. 43, 932–44 (2010).

9. Biton, A. et al. Independent component analysis uncovers the landscape of the bladder tumor transcriptome and reveals insights into luminal and basal subtypes. Cell Rep. 9, 1235–45 (2014).

10. Van Der Maaten, L. & Hinton, G. Visualizing Data using t-SNE. J. Mach. Learn. Res. 9, 2579–2605 (2008).

11. Subramanian, A. et al. Gene set enrichment analysis: a knowledge-based approach for interpreting genome-wide expression profiles. Proc. Natl. Acad. Sci. U. S. A. 102, 15545–50 (2005).

12. Lim, S. et al. Lysine-specific demethylase 1 (LSD1) is highly expressed in ER-negative breast cancers and a biomarker predicting aggressive biology. Carcinogenesis 31, 512–520 (2010).

13. Nagasawa, S. et al. LSD1 Overexpression Is Associated with Poor Prognosis in Basal-Like Breast Cancer, and Sensitivity to PARP Inhibition. PLoS One 10, e0118002 (2015).

14. Himberg, J., Hyvärinen, A. & Esposito, F. Validating the independent components of neuroimaging time series via clustering and visualization. Neuroimage 22, 1214–1222 (2004).

15. Jahirabadkar, S. & Kulkarni, P. Clustering for High Dimensional Data: Density based Subspace Clustering Algorithms. Int. J. Comput. Appl. 63, 975–8887 (2013).

16. Koboldt, D. C. et al. Comprehensive molecular portraits of human breast tumours. Nature 490, 61–70 (2012).

17. Hyvärinen, A. Independent component analysis: recent advances. Philos. Trans. R. Soc. London A Math. Phys. Eng. Sci. 371, (2012).

18. Hyvärinen, A. & Oja, E. Independent component analysis: algorithms and applications. Neural Networks 13, 411–430 (2000).

19. Mootha, V. K. et al. PGC-1α-responsive genes involved in oxidative phosphorylation are coordinately downregulated in human diabetes. Nat. Genet. 34, 267–273 (2003).

20. Shannon, P. et al. Cytoscape: A Software Environment for Integrated Models of Biomolecular Interaction Networks. Genome Res. 13, 2498–2504 (2003).

